# Latent class regression improves the predictive acuity and clinical utility of survival prognostication amongst chronic heart failure patients

**DOI:** 10.1101/2020.11.27.400887

**Authors:** John L Mbotwa, Marc de Kamps, Paul D Baxter, George TH Ellison, Mark S Gilthorpe

## Abstract

The present study aimed to compare the predictive acuity of latent class regression (LCR) modelling with: standard generalised linear modelling (GLM); and GLMs that include the membership of subgroups/classes (identified through prior latent class analysis; LCA) as alternative or additional candidate predictors. Using real world demographic and clinical data from 1,802 heart failure patients enrolled in the UK-HEART2 cohort, the study found that univariable GLMs using LCA-generated subgroup/class membership as the sole candidate predictor of survival were inferior to standard multivariable GLMs using the same four covariates as those used in the LCA. The inclusion of the LCA subgroup/class membership together with these four covariates as candidate predictors in a multivariable GLM showed no improvement in predictive acuity. In contrast, LCR modelling resulted in a 10-14% improvement in predictive acuity and provided a range of alternative models from which it would be possible to balance predictive acuity against entropy to select models that were optimally suited to improve the efficient allocation of clinical resources to address the differential risk of the outcome (in this instance, survival). These findings provide proof-of-principle that LCR modelling can improve the predictive acuity of GLMs and enhance the clinical utility of their predictions. These improvements warrant further attention and exploration, including the use of alternative techniques (including machine learning algorithms) that are also capable of generating latent class structure while determining outcome predictions, particularly for use with large and routinely collected clinical datasets, and with binary, count and continuous variables.

## Introduction

### The limited acuity and clinical utility of Generalised Linear Models (GLMs)

The potential utility of predictive modelling, using routinely collected data for diagnosis, prognostication and health service planning, is one of five ‘novel capabilities’ that Wang et al. [1] identified as pertinent to the application of data analytics in medicine and health. Long before John Mashey first applied the term ‘Big Data’ in this context during the late 1990s [2], generalised linear models (GLMs) were used to develop clinical ‘risk scores’ based on much smaller scale datasets [3]. Indeed, clinical prediction models (CPMs) remain popular for prognostication and are in widespread usage to this day, not least in cardiovascular medicine [4,5].

While CPMs and their wider utility remain contentious (beyond strict prognostication, and particularly in prevention [6-9]), many of the standard statistical modelling techniques commonly used are on clinical datasets that remain relatively small – at least when compared to contemporary notions of ‘Big Data’ [2]. A substantial statistical weakness of the commonest of these (generalised linear models; GLMs) as a predictive tool is that they often fail to make *full* use of the joint information available amongst *all* candidate predictor variables. This is because these models rarely explore nonlinear relationships and interactions. Moreover, even when analysts optimally parameterise the candidate predictors available, and carefully consider all possible interaction terms between these, the clinical utility of GLMs is typically limited to predictions made at the population level [6,10], while predictions at the individual level often lack precision (and with it, utility).

Although more sophisticated machine learning techniques may overcome the rigidity of GLMs and analysts’ tendency to ignore (or overlook) nonlinear relationships and interactions, population-level predictions generated using cutting edge machine learning techniques will still be more reliable than individual-level predictions. Indeed, this bald fact applies to all prediction modelling techniques, including those underpinning contemporary claims of ‘personalised’ or ‘precision medicine’ [11]. It is therefore critical to recognise that while it is possible to determine what proportion of any given population will experience a specified outcome with a reasonable degree of accuracy, all such models provide less accuracy in determining outcomes for each *individual* within that population.

Meanwhile, a further epistemological consideration that commonly arises in CPMs (and elsewhere) is the mistaken belief that the coefficient estimates of covariates included/retained in the model indicate the extent to which each covariate contributes to the model’s overall prediction (of the model’s specified outcome). This belief is mistaken because each covariate’s coefficient estimate is generated *conditional* on the adjustment of all other covariates in the model, such that the contribution of any one covariate is merged with that of all other covariates included in that model. For this reason, what the coefficient estimate of each covariate actually represents is the residual relationship between that covariate and the outcome *subject to* the joint contributions made by all other covariates in the model *considered simultaneously*.

This situation is further complicated where any of the covariates included in the model reflect events that occurred contemporaneously with or even after the specified outcome – in this instance, the joint contributions made by all covariates would be subject to conditioning on the outcome, which can have other adverse consequences on model interpretation [12-14]. In practice, the inclusion of covariates acting contemporaneously with or after the outcome’ in prediction modelling is likely to be used only where the aim is to estimate the values for variables whose measurements are missing, imprecise or challenging to measure (i.e. in modelling that aims to achieve what might be called ‘predictive interpolation’ for diagnostic and related measurement/ascertainment purposes). These issues aside, it is important to stress that the coefficient estimates of all covariates (with the exception of the covariate closest in time to the outcome) cannot be causally interpreted, as they will be subject to inferential bias known as mediator bias [15], which undermines causal interpretation of their coefficients due to the so-called ‘Table 2 Fallacy’ [16].

Due to these caveats, predictions that are generated using GLMs cannot address the two key concerns of attending physicians, namely: “Which of the covariates (i.e. ‘predictor’ variables) are amenable to clinical intervention, so as to prevent or mitigate any adverse outcome (or promote and amplify any favourable outcome) in each (or all) of these patients?” and “Which particular patients will experience an adverse (or favourable) outcome?”. To address the first of these questions, analysts need to switch their focus from predicting outcome values to estimating each of the relationships between covariates considered plausible targets for intervention and the outcome – an approach that can capitalise on recent advances in causal inference modelling techniques [17]. To address the second question, the best that can be achieved is to identify clinically meaningful *subgroups* of patients with shared characteristics that set them apart from other (*subgroups* of) patients – a relatively novel approach that involves multivariable ‘risk profiling’.

### Improving the acuity and clinical utility of predictive modelling for prognostication

Multivariable risk profiling can be achieved using latent class analysis (LCA) in which the exploration of nonlinearity, and of important interactions amongst included covariates, forms an integral (albeit implicit) part of classifying patients into subgroups [18]. Despite these benefits, the clinical utility of the resulting latent classes ultimately depends upon the extent to which this approach optimally exploits the joint information amongst available covariates. This approach perhaps has greatest clinical utility where there are: (i) factors known to be *associated* with the outcome (which therefore facilitate prediction); but (ii) there are no known, modifiable causes of the outcome, or aetiological understanding is poor/contested (as is the case with many rare, novel or complex diseases). Indeed, providing that the specified outcome is excluded from the LCA process (to avoid conditioning on the outcome) [14], combining LCA class membership with candidate predictors provides increased complexity that can help exploit the joint covariate information in multivariable GLM prediction. That said, it is important to stress that causal interpretation of any covariate coefficients for latent class membership in such models remains deeply flawed for the very same reasons that causal interpretations of any covariate coefficient in prediction GLMs is flawed (as explained earlier). Ostensibly this consideration might appear to limit the clinical utility of LCA-generated class membership, and it is true that describing class membership as a ‘risk factor’ often generates, and commonly reflects, a lack of understanding. Indeed, it risks conflating prediction and causal inference/determination just as it does when individual covariates are described in similar terms as ‘risk factors’ [7].

Thus, while classifying subgroups of individuals using LCA can improve analytical practice and strengthen consideration of nonlinear relationships (and important interactions amongst covariates), it does not address the clinical appetite for identifying so-called ‘modifiable risk factors’, or for individually tailored risk probabilities (the so-called ‘holy grail’ of personalised or precision medicine) [10]. This might explain why the use of latent variable methods in prediction modelling remains largely under-explored, even though more sophisticated approaches exist that incorporate such techniques within GLM and offer substantial advantages for clinicians through subgroup risk profiling. These approaches involve the construction of latent classes ‘across’ multivariable GLMs to: integrate consideration of nonlinear relationships and important interactions between covariates; and better capture (and exploit) the joint information amongst the available/included covariates. For example, in what is termed latent class regression (LCR) modelling, population data are partitioned into their constituent latent classes and a distinct GLM is simultaneously generated for each class. In the process, this approach accommodates any inherent population heterogeneity and thereby improves model precision.

In its simplest form, LCR models may be viewed as two distinct modelling concepts undertaken in a single estimation process: in the first, population data are probabilistically assigned to latent classes (population subgroups); while, in the second, separate GLMs are derived for each class/subgroup. The probability of an individual belonging to each class is based on similarities in the characteristics displayed by individuals attributed to different classes. Importantly, the assignment of individuals to classes is not limited to just those covariates available for analysis, since outcome differences attributable to unknown (i.e. latent) influences are also accommodated, *and* without inappropriate conditioning on the outcome. Individuals may thus belong to more than one class, with the sum of probabilities over all classes being one. Within each class, distinct GLMs are generated, with the selection of covariates acting as predictors (and their model coefficients) permitted to vary from one class to the next. In this way, by ensuring that the consideration of potential nonlinearity and possible interactions is integral to the application of LCR models, these models should exploit the covariate joint information available in a more consistent fashion and thereby strengthen the acuity of the prediction achieved. An additional benefit of this approach is that the latent classes/subgroups identified using LCR may also strengthen the clinical utility of the prediction achieved because any variation in the risk of the outcome amongst different classes/subgroups can be used to target diagnostic, therapeutic or palliative resources more precisely and efficiently.

The aim of the present study was therefore to explore whether LCR models might improve the accuracy and precision of predictions at the population *and* individual level, by comparing LCR-generated predictions to standard GLM and LCA-informed GLM (including the use of LCA-generated class membership as either the *only* candidate predictor in univariable GLMs, or as an *additional* candidate predictor alongside all other available covariates in multivariable GLMs). We thus explore four models offering progressively more complex exploitation of the individual and joint information available from the covariates available for consideration as candidate predictors. To this end, the analyses that follow use data (on age, sex, haemoglobin level and diabetes) that are routinely available in a clinical context (cardiovascular medicine) in which Cox proportional hazards time-to-event analyses are commonly used in prognostic predictions of mortality, where survival and loss to follow-up are pertinent analytical endpoints.

## Methods

### Study design, data collection and ethics

The analyses that follow used data from the United Kingdom Heart Failure Evaluation and Assessment of Risk Trial 2 (UK-HEART2) – a prospective cohort of ambulant patients with signs and symptoms of chronic heart failure (CHF) [19]. The study recruited 1,802 adult patients with CHF who attended specialist cardiology clinics in four UK hospitals between July 2006 and December 2014 [20]. Patients were eligible for recruitment if they: were aged 18 years or older; had had clinical signs and symptoms of CHF for at least 3 months; and had a left ventricular ejection fraction that was less than or equal to 45% [19,20]. Ethical approval was obtained from the research ethics committee at each participating hospital and eligible study participants were only recruited following informed consent [21]. Additional information regarding UK-HEART-2’s study design, patient eligibility and inclusion criteria, together with a detailed description of the study cohort has been reported elsewhere [19-21].

#### Statistical methods

To simplify the methodological comparisons undertaken in the present study, the covariates selected as candidate predictors comprised two demographic variables (age, sex), a single physiological parameter (haemoglobin level), and a single clinical characteristic (type 2 diabetes). These four covariates were then used to generate prognostic predictions of survival amongst UK-HEART-2 participants using four separate statistical Procedures (for each of which the underlying principles and model building processes are described in detail in Part 1 of the Supplementary Materials):

- Procedure 1 (standard GLM) involved single step multivariable Cox proportional hazards models that considered all four covariates as candidate predictors of survival, with no consideration of nonlinear relationships or interactions between covariates.
- Procedure 2 (LCA-informed GLM *without* the inclusion of covariates) involved two sets of models, each involving two separate steps. First, LCA was used to identify any latent classes/subgroups of participants using the four selected covariates, with individual membership to each latent class allocated using modal (Procedure 2a) and probabilistic (Procedure 2b) assignment. Second, *univariable* Cox proportional hazards models examined latent class membership as the *sole* predictor of survival, with separate models generated using latent class membership derived using modal (Procedure 2a) or probabilistic (Procedure 2b) assignment.
- Procedure 3 (LCA-informed GLM *with* the inclusion of covariates) again involved two sets of models, each involving two separate steps. First, LCA was used to allocate latent class membership using modal (Procedure 3a) and probabilistic (Procedure 3b) assignment – as in the first step of Procedure 2 (above). Second, *multivariable* Cox proportional hazards models considered all four covariates (as used in Procedure 1) *plus* latent class membership as *multiple* predictors of survival, with separate models generated using latent class membership derived using probabilistic (Procedure 3a) or modal (Procedure 3b) assignment.
- Procedure 4 (LCR) involved single step latent class regression (LCR) models that considered all four covariates as candidate predictors to simultaneously predict *both* latent class membership *and* survival within each latent class.

For all latent class models, entropy is reported which assesses the extent that individuals are aligned predominantly to a single class (i.e. having a large modal probability, leading to a greater entropy), as this facilitates clearer interpretation of each latent class as a near-complete collection of individuals. Model optimisation in terms of the number of latent classes may thus depend upon *both* the overall predictive acuity of the latent class structure (as evident from the model BIC) *and* the intended utility of the determined classes thereafter (as indicated by the model entropy). For this illustration, we prioritise overall predictive acuity.

All descriptive statistics and GLMs were generated using *R* (version 4.0.3) [22], as were the model specification, selection, validation and bootstrapping procedures (Part 2 of the Supplementary Materials). All latent class modelling was undertaken in M*plus* (version 8.3) [23], using the M*plus* automation package to run models in M*plus* from within *R* [22].

## Results

The first column of Table 1 summarises the distribution of each covariate amongst participants in the UK-HEART-2 cohort. These indicate that: the mean age of the cohort’s participants was 70 years; around two thirds (69.7%) were male; over a quarter (28%) had type 2 diabetes; the mean level of circulating haemoglobin was 13.5 g/dl; and 60% died during the period of follow-up (equivalent to a median survival of 3.4 years).

**Table 1.**
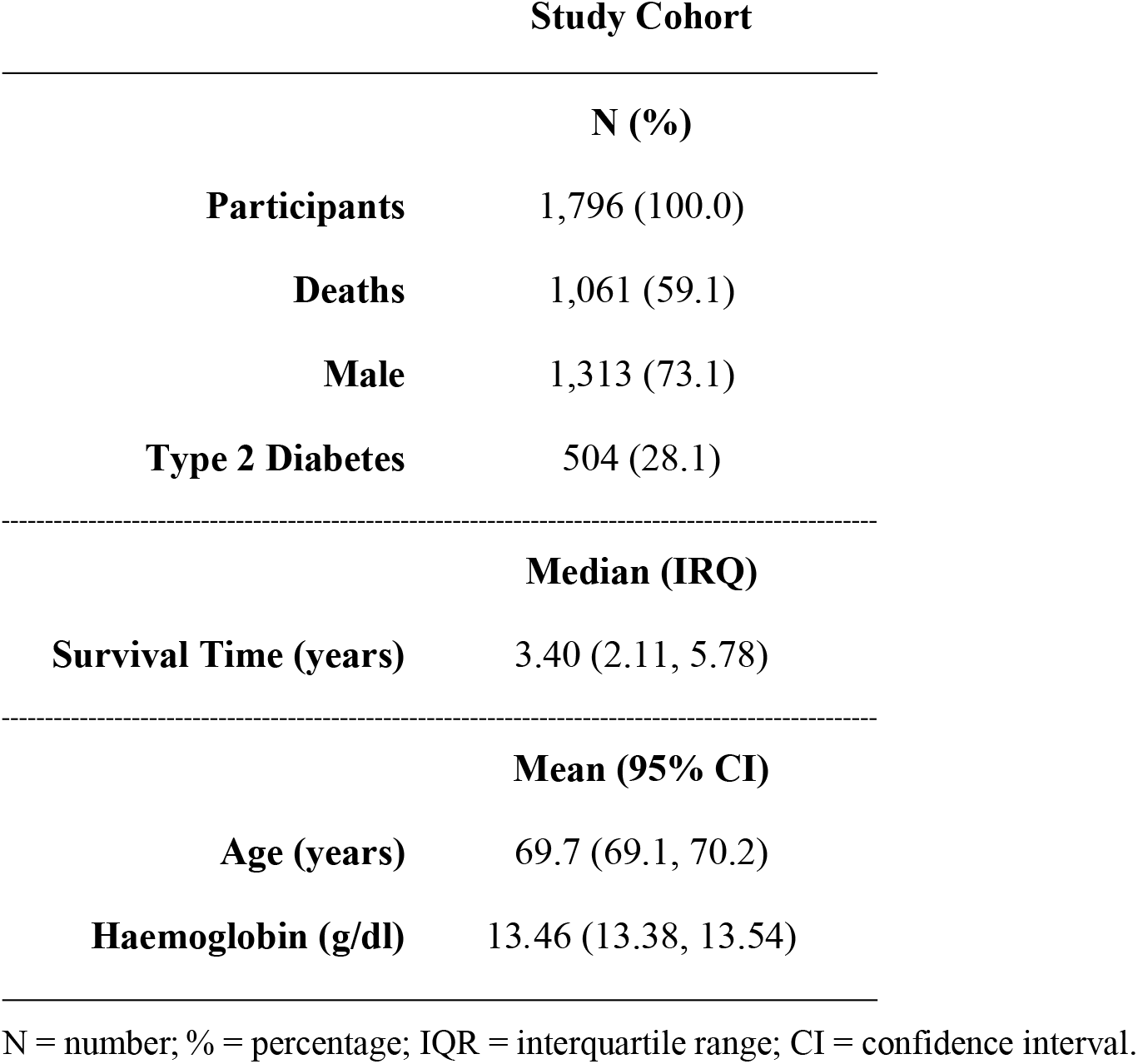
Descriptive characteristics of the study cohort.

**Table 2.** Latent class analysis (LCA) model summaries – the preferred model from this step was used in Procedures 2 and 3.

| Number of classes | Number of parameters | BIC | Entropy | Class | Modal N (%) | Probabilistic N (%) |
| --- | --- | --- | --- | --- | --- | --- |
| 1 | 6 | 19,818.53 | - |  | 1,796 (100.0) | - |
| 2 | 11 | 19,537.79 | 0.75 | Class 1 | 1,452 (80.8) | 1425.3 (79.4) |
|  |  |  |  | Class 2 | 344 (19.2) | 370.7 (20.6) |
| 3 | 16 | 19,445.74 | 0.74 | Class 1 | 1,203 (67.0) | 11744.0 (65.4) |
|  |  |  |  | Class 2 | 480 (26.7) | 500.7 (27.9) |
|  |  |  |  | Class 3 | 113 (6.3) | 120.3 (6.7) |
| 4 | 21 | 19,422.35 | 0.80 | Class 1 | 811 (45.2) | 797.0 (44.4) |
|  |  |  |  | Class 2 | 486 (27.1) | 504.4 (28.1) |
|  |  |  |  | Class 3 | 381 (21.2) | 371.4 (20.7) |
|  |  |  |  | Class 4 | 118 (6.6) | 123.2 (6.9) |
| 5 | 26 | 19,421.44 | 0.67 | **Class 1** | **586 (32.6)** | **566.7 (31.6)** |
|  |  |  |  | **Class 2** | **470 (26.2)** | **459.7 (25.6)** |
|  |  |  |  | **Class 3** | **324 (18.0)** | **296.9 (16.5)** |
|  |  |  |  | **Class 4** | **317 (17.7)** | **368.6 (20.5)** |
|  |  |  |  | **Class 5** | **99 (5.5)** | **104.1 (5.8)** |
| 6 | 31 | 19,422.87 | 0.63 | Class 1 | 527 (29.3) | 517.7 (28.8) |
|  |  |  |  | Class 2 | 474 (26.4) | 470.5 (26.2) |
|  |  |  |  | Class 3 | 276 (15.4) | 247.7 (13.8) |
|  |  |  |  | Class 4 | 234 (13.0) | 232.6 (13.0) |
|  |  |  |  | Class 5 | 186 (10.4) | 229.8 (12.8) |
|  |  |  |  | Class 6 | 99 (5.5) | 97.6 (5.4) |

BIC = Bayesian information criterion; N = number; % = percentage; the optimal LCA model according to the BIC is emboldened.

In Procedure 1, the single step Cox proportional hazards models that considered all four covariates as candidate predictors of survival found that the model in which all four covariates were retained achieved the highest AUC (0.69) – a level of acuity considered ‘modest to poor’ [24].

In Procedure 2, the LCAs conducted during the first step found that the 5-class model which retained all four covariates had the most favourable BIC (Table 2). Applying this 5-class model during the second step as the sole predictor of survival in a Cox proportional hazards model, achieved an AUC of 0.65 using modal assignment (Procedure 2a) and 0.66 using probabilistic assignment (Procedure 2b). These levels of acuity were both lower than that achieved using Procedure 1 (AUC=0.69).

In Procedure 3, the second step involved consideration not only of the four covariates as candidate predictors of survival in the Cox proportional hazards model (as in Procedure 1), but also membership of the same 5-class model developed in the first step of Procedure 2. These analyses found that: the best fitting GLMs did not retain class membership as a predictor; and forcibly retaining class membership in the model did not improve the AUC above that achieved in Procedure 1 or 2, regardless of how class membership was assigned (modal: AUC=0.65; probabilistic: AUC=0.66).

In Procedure 4, with all four covariates eligible for inclusion as candidate predictors of *both* latent class membership *and* the Cox Proportional Hazards models, some of the models were over-parameterised and failed to converge. Nonetheless, the most favourable of the models that successfully converged involved a latent class variable with just two classes and an AUC of 0.79 (Table 3). When compared to the best performing models in Procedures 1-3, these results suggest that Procedure 4 achieved a substantial improvement in predictive acuity of 10-14%.

**Table 3.** Latent class regression (LCR) model summaries for Procedure 4.

| Number<br>of classes | Number of<br>parameters | BIC | Entropy |
| --- | --- | --- | --- |
| 1 | 3 | 3695.06 | ---- |
| **2** | **10** | **3659.49** | **0.68** |
| 3 | 17 | 3682.44 | 0.91 |
| 4 | 24 | 3722.89 | 0.94 |

BIC = Bayesian information criterion; the optimal LCA model according to the BIC is emboldened.

Improvements in acuity aside, the most favourable of the LCR models had only three of the covariates (age, sex, and type 2 diabetes) retained in the Cox proportional hazards models for each membership class, and only one of these covariates (type 2 diabetes) and the remaining covariate (haemoglobin level) retained as covariates in the LCR class membership model (Table 4). Given that all four covariates were retained in the most favourable CPH models generated by Procedures 1 and 3, and in the LCA models generated in the first step of Procedures 2 and 3, these findings suggest that Procedure 4’s 10-14% improvement in AUC is likely to have been achieved by exploiting the available covariate information differently to each of the three other Procedures. An indication of what this entailed can be found in the distribution of covariate characteristics amongst the two classes of the most favourable LCR model (Table 5), which suggest that these classes might warrant post-hoc labelling as ‘high risk’ and ‘low risk’ subgroups and might thereby offer substantial additional clinical utility (in guiding the allocation of diagnostic, therapeutic and/or palliative resources).

**Table 4.**
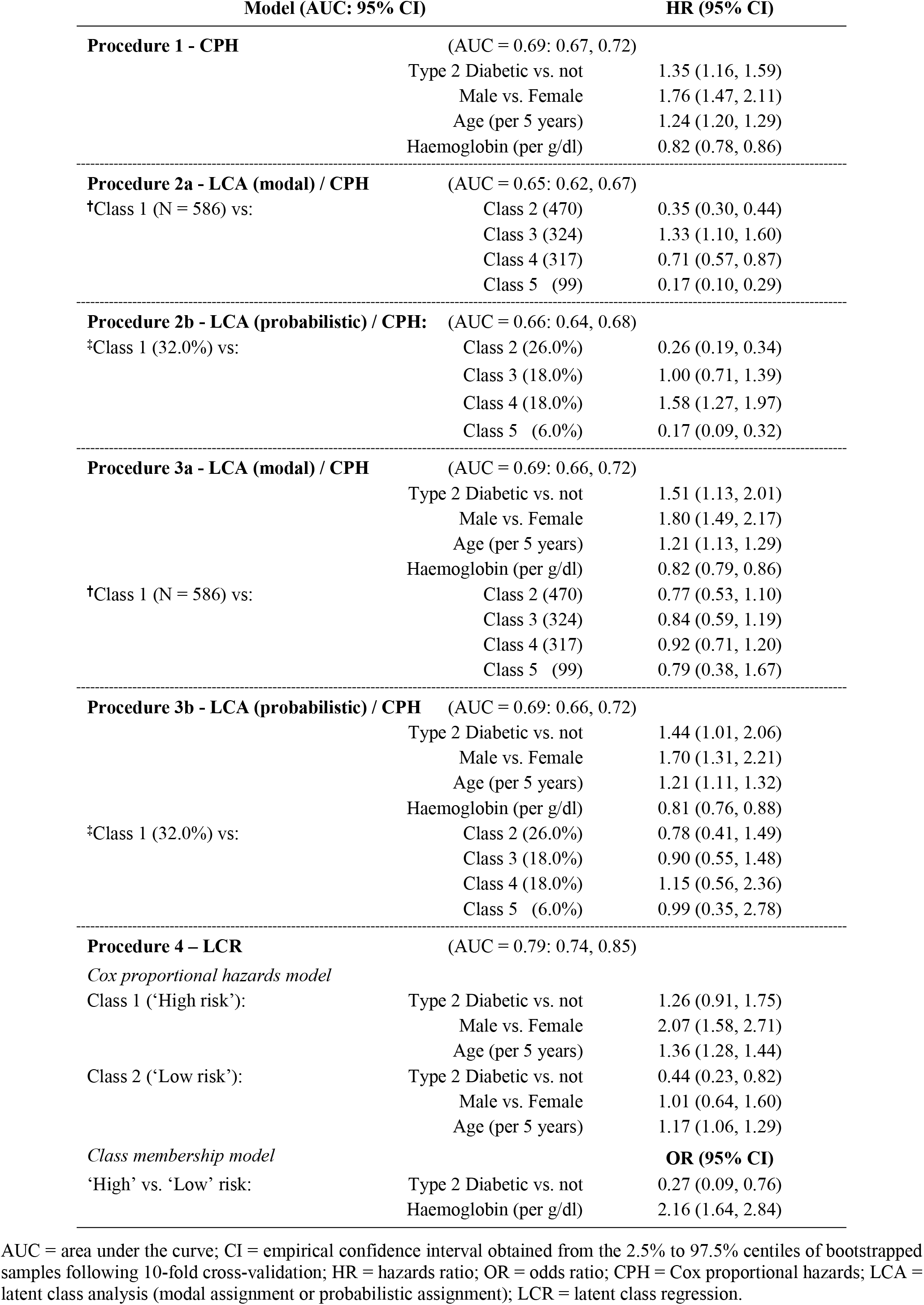
Covariate coefficients for each preferred model (Procedures 1-4) executed on the complete data, along with median AUC and empirical 95% empirical confidence intervals generated through 10-fold cross-validation.

| Model (AUC: 95% CI) |  | HR (95% CI) |
| --- | --- | --- |
| **Procedure 1 - CPH** (AUC = 0.69: 0.67, 0.72) |  |  |
|  | Type 2 Diabetic vs. not | 1.35 (1.16, 1.59) |
|  | Male vs. Female | 1.76 (1.47, 2.11) |
|  | Age (per 5 years) | 1.24 (1.20, 1.29) |
|  | Haemoglobin (per g/dl) | 0.82 (0.78, 0.86) |
| **Procedure 2a - LCA (modal) / CPH** (AUC = 0.65: 0.62, 0.67) |  |  |
| †Class 1 (N = 586) vs: | Class 2 (470) | 0.35 (0.30, 0.44) |
|  | Class 3 (324) | 1.33 (1.10, 1.60) |
|  | Class 4 (317) | 0.71 (0.57, 0.87) |
|  | Class 5 (99) | 0.17 (0.10, 0.29) |
| **Procedure 2b - LCA (probabilistic) / CPH:** (AUC = 0.66: 0.64, 0.68) |  |  |
| ‡Class 1 (32.0%) vs: | Class 2 (26.0%) | 0.26 (0.19, 0.34) |
|  | Class 3 (18.0%) | 1.00 (0.71, 1.39) |
|  | Class 4 (18.0%) | 1.58 (1.27, 1.97) |
|  | Class 5 (6.0%) | 0.17 (0.09, 0.32) |
| **Procedure 3a - LCA (modal) / CPH** (AUC = 0.69: 0.66, 0.72) |  |  |
|  | Type 2 Diabetic vs. not | 1.51 (1.13, 2.01) |
|  | Male vs. Female | 1.80 (1.49, 2.17) |
|  | Age (per 5 years) | 1.21 (1.13, 1.29) |
|  | Haemoglobin (per g/dl) | 0.82 (0.79, 0.86) |
| †Class 1 (N = 586) vs: | Class 2 (470) | 0.77 (0.53, 1.10) |
|  | Class 3 (324) | 0.84 (0.59, 1.19) |
|  | Class 4 (317) | 0.92 (0.71, 1.20) |
|  | Class 5 (99) | 0.79 (0.38, 1.67) |
| **Procedure 3b - LCA (probabilistic) / CPH** (AUC = 0.69: 0.66, 0.72) |  |  |
|  | Type 2 Diabetic vs. not | 1.44 (1.01, 2.06) |
|  | Male vs. Female | 1.70 (1.31, 2.21) |
|  | Age (per 5 years) | 1.21 (1.11, 1.32) |
|  | Haemoglobin (per g/dl) | 0.81 (0.76, 0.88) |
| ‡Class 1 (32.0%) vs: | Class 2 (26.0%) | 0.78 (0.41, 1.49) |
|  | Class 3 (18.0%) | 0.90 (0.55, 1.48) |
|  | Class 4 (18.0%) | 1.15 (0.56, 2.36) |
|  | Class 5 (6.0%) | 0.99 (0.35, 2.78) |
| **Procedure 4 – LCR** (AUC = 0.79: 0.74, 0.85) |  |  |
| *Cox proportional hazards model* |  |  |
| Class 1 ('High risk'): | Type 2 Diabetic vs. not | 1.26 (0.91, 1.75) |
|  | Male vs. Female | 2.07 (1.58, 2.71) |
|  | Age (per 5 years) | 1.36 (1.28, 1.44) |
| Class 2 ('Low risk'): | Type 2 Diabetic vs. not | 0.44 (0.23, 0.82) |
|  | Male vs. Female | 1.01 (0.64, 1.60) |
|  | Age (per 5 years) | 1.17 (1.06, 1.29) |
| *Class membership model* |  | **OR (95% CI)** |
| 'High' vs. 'Low' risk: | Type 2 Diabetic vs. not | 0.27 (0.09, 0.76) |
|  | Haemoglobin (per g/dl) | 2.16 (1.64, 2.84) |

AUC = area under the curve; CI = empirical confidence interval obtained from the 2.5% to 97.5% centiles of bootstrapped samples following 10-fold cross-validation; HR = hazards ratio; OR = odds ratio; CPH = Cox proportional hazards; LCA = latent class analysis (modal assignment or probabilistic assignment); LCR = latent class regression.

**Table 5.** Descriptive characteristics for the 2-class Cox proportional hazards latent class regression model.

|  | Latent Class Regression Model |  |  |  |
| --- | --- | --- | --- | --- |
|  | Class 1 ('High risk') |  | Class 2 ('Low risk') |  |
|  | Modal N (%) | Probabilistic N (%) | Modal N (%) | Probabilistic N (%) |
| Participants | 1,566 (87.2) | 1507.8 (84.0) | 230 (22.8) | 288.2 (16.0) |
| Deaths | 1,046 (66.8) | 1014.7 (67.3) | 15 (6.5) | 45.8 (15.9) |
| Male | 1,160 (74.1) | 1112.8 (73.8) | 153 (66.5) | 200.9 (69.7) |
| Type 2 Diabetes | 368 (23.5) | 342.3 (22.7) | 136 (59.1) | 162.5 (56.4) |
|  | Median (IRQ) |  | Median (IRQ) |  |
| Survival Time (years) | 3.86 (2.41, 5.89) |  | 1.13 (0.50, 2.27) |  |
|  | Mean (95% CI) |  | Mean (95% CI) |  |
| Age (years) | 69.2 (68.6, 69.9) |  | 72.5 (71.1, 73.9) |  |
| Haemoglobin (g/dl) | 13.80 (13.72, 13.88) |  | 11.14 (10.99, 11.30) |  |

N = number; % = percentage; IQR = interquartile range; CI = confidence interval.

A further key finding that emerges from closer examination of the Cox proportional hazards models generated for each of the two classes within the optimum LCR model (Table 4) is that the contribution made by each of the covariates therein varied by class, and was dissimilar to the contribution these covariates made in those Procedures where all covariates were available for inclusion as separate candidate predictors (i.e. Procedure 1 and 3a/b). While the coefficient estimates of covariates in each of these models cannot be interpreted as measures of causal effects [16], their contribution as candidate predictors is strikingly different and depends upon the choice of model(s) used in each Procedure (Table 4). For example, the hazard of death associated with being male was 1.7 to 1.8 in Procedures 1 and 3, whilst for Procedure 4 being male was associated with a substantially higher hazard of death in one class (HR = 2.07; 1.58, 2.71) yet was unrelated to the hazard of death in the other class (HR = 1.01; 0.64, 1.60). Likewise, Type 2 diabetes was consistently associated with an elevated hazard of death in models generated under Procedure 1 and 3, while in Procedure 4 this covariate was associated with both an *elevated* hazard of death in one class (HR = 1.26; 0.91, 1.75) and a *reduced* hazard of death in the other class (HR = 0.43; 0.23, 0.82).

Clearly, the joint information available amongst each of the candidate predictors is selected and utilised very differently by each of the Procedures examined in the present study (see Table 4). Nonetheless, what sets the LCR model in Procedure 4 apart from the models used in Procedures 1-3 is that LCR allows the predictive contribution from each covariate to be partitioned *across* any latent substructures existing within the study population, such that covariates are able to operate differently within each of the latent subgroups – thereby capturing and reflecting population heterogeneity that is: unavailable to any of the other modelling Procedures; and, crucially, of substantial (additional) value when predicting the specified outcome.

## Discussion

The present study provides proof of principle that LCR models can provide substantive improvements in predictive acuity *and* clinical utility over standard approaches using GLM (with or without LCA). Nonetheless, there are several potential limitations that warrant consideration and further investigation. In particular, it would be insightful to compare these alternative approaches to prediction using larger datasets and larger numbers of covariates than those chosen in this instance for illustration. This might involve comparing Procedures 1 through 4 using different numbers and sets of covariates from similar sized datasets; as well as extending the application of LCR modelling to more complex scenarios and much larger datasets. At the same time, it is important to point out that, in the context of the dataset used in the present study, the underlying ‘truth’ (and the data generating mechanisms involved) cannot be known with certainty, and exploring the potential strengths (and analytical limitations) of LCR would thus benefit from extensive simulations to evaluate a range of different circumstances for a range of different covariates *and* outcomes (including those comprising binary, continuous and count variables) to evaluate whether LCR continues to perform well (and better than GLM, LCA or both) under these circumstances. In the absence of subsequent research along these lines, the ‘proof of principle’ offered by the present study remains speculative; although it would also be worth exploring whether alternative approaches to prediction modelling might be incorporated into, or integrated with, the analytical principles underpinning LCR modelling, such as the inclusion of similar dual modelling structures within machine learning, to assess whether the apparent benefits of LCR models might be further enhanced.

These limitations, the present study successfully compared three different approaches for incorporating latent variable methods within prediction modelling and demonstrated that LCR models can outperform not only the standard approach using GLM (in which membership of latent classes is ignored – Procedure 1), but also those that include latent class membership identified using LCA to generate an alternative (Procedure 2) or additional (Procedure 3) candidate predictor. This improvement in predictive acuity (which, as shown above, resulted in a 10-14 percentage point improvement in AUC, despite the modest number of participants and covariates involved) illustrates the potential benefits of LCR for prediction modelling which, in this instance, shifted the acuity of prediction from ‘modest to poor’ to ‘substantial’ [24].

The present study also demonstrated that the latent class/subgroup structure that is revealed through LCR may have potential clinical utility. This is because it might – as in the example examined here – facilitate the identification of discrete *sub*groups (i.e. latent classes) of populations with very different underlying risks of the outcome. While such subgroups may not necessarily be amenable to effective intervention (given that LCR models support prediction, not causal inference [16]), they should help to improve the efficient allocation/targeting of outcome-relevant diagnostic, therapeutic and/or palliative resources to those subgroups identified as more likely to require (and perhaps even benefit from) these. However, to maximise the clinical exploitation of latent subgroups identified using LCR (and similar techniques), model selection must focus on those achieving higher entropy – where the probability of class assignment is closer to one for most assignments – as this better aligns individuals/participants to a predominant single class (rather than aligning individuals/participants to multiple classes). For example, in Procedure 4, the 3-class model had lower predictive acuity but greater entropy than the 2-class model (see Table 3); and had the identification of clinically meaningful subgroups been the focus of these analyses (as opposed to overall predictive acuity), then it might have been appropriate to accept a modest reduction in predictive acuity in favour of enhanced clinical utility – i.e. recognising three (‘high’, ‘medium’ and ‘low’ risk) subgroups rather than just the two (‘high’ and ‘low’ risk) subgroups identified by the LCR model with the most favourable predictive acuity (Table 3). Indeed, when clinical resources are scarce, such an approach might prove a more reliable approach to resource allocation than one based upon a stringent interpretation of predictive acuity alone.

## Online Supplementary Material

### Part 1. Underlying principles and model-building processes involved in the GLM, LCA and LCR techniques examined

#### GLM: the Cox proportional hazards model

A Cox proportional hazards model generates a (hazard) function which indicates the risk of the outcome occurring during the period of follow-up. Mathematically, a Cox regression model [25,26] is defined as:

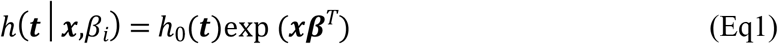

where: ***t*** is a non-negative random variable representing time to ‘death’, ‘loss to follow-up’ or ‘the end of the study’ for all participants (in this example, patients with CHF); ℎ_0_(***t***) is the baseline hazard function; ***x*** is the vector of predictors for the time-to-event outcome ***t***; and ***β**^T^* is the transpose of the vector of coefficients obtained from the Cox proportional hazards model. To make predictions using the Cox proportional hazards model, the survival function is defined as:

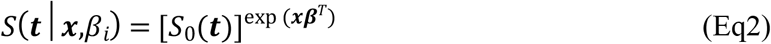

where, if the baseline hazard function ℎ_0_(***t***) is known, then:

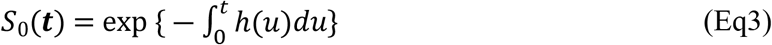

#### LCA: the general latent class (profile) model

Latent class (profile) models come from a family of finite mixture models that classify observations into classes associated with unobserved heterogeneity in a population. A population is partitioned into *g* classes for the outcome ***y*** with the mixture density function defined in relation to covariates ***x*** as:

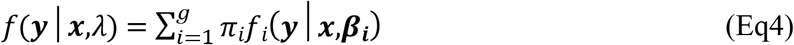

where *f_i_*(***y***|***x***,***β_i_)*** is the conditional probability density function for the observed response in the *i^th^* class and π_***i***_ (*i* = 1…*g*) represent the class-membership probabilities that are estimated for each class such that:

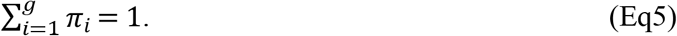

For a class membership model, the structural part of the model is given by:

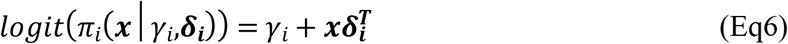

hence

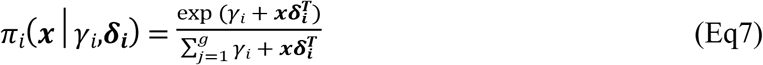

where: ***x*** is a (*p* × 1) covariate vector for the class-membership model; and 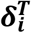 is the transposition of the vector ***δ_i_*** for the multinomial logistic class-membership model.

#### LCR: the latent class regression model

The latent class regression (LCR) model is an extended version of the generalised linear model where the concept of latent class mixtures is applied to the entire model specified, not just to a cluster of covariates. LCR survival analysis extends this to the time-to-event framework of Cox proportional hazards modelling to: (i) predict probabilistically assigned subgroups of participants with different futures (in this example, subgroups of patients with different prognoses of survival/death) based on the available covariates; and, (ii) *simultaneously* predict the survival distributions for each subgroup selecting from the same covariates acting as candidate predictors. The distribution of the survival time variable for each component in Eq4 can be:

- parametric – a scenario *with* distributional assumptions concerning the survival times;
- semi-parametric – a scenario with *relaxed* distributional assumptions; or
- non-parametric – a scenario *without* distribution assumptions concerning the survival times.

Assuming a parametric model for the specified outcome variable, the component’s densities are assumed to be from the same family, so that a number of common distribution functions may be considered appropriate for survival times, such as: exponential; Gamma; and Weibull [38]. In a semi-parametric case, the Cox proportional hazards model is an example. Within a latent class framework, if ***t*** is the random variable representing time to event (e.g. ‘death’, ‘loss to follow-up’, or ‘end of the study’), individuals are divided into *g* latent classes that are differentiated by the covariate vector ***x***, with individual survival in each class *i* predicted by covariate vector ***x***, and the survival model is defined as:

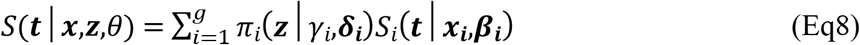

where: *θ* = (*γ*_*i*_, ***δ_i_***, ***β_i_***) is the collection of parameters to be estimated such that *π*_*i*_(***z***|*γ*_*i*_,***δ_i_***) satisfies the constraints in Eq4. Vectors ***x_i_*** and ***z*** are any available measures of participant characteristics, exposures and treatments etc., which may be the same or differ, just as the ***x_i_*** covariates may also differ for each class.

If the effects of the ***x_i_*** covariates on the hazards (i.e. the instantaneous risk of event) in each class is constant during the duration of follow-up, then the hazard function can be specified as:

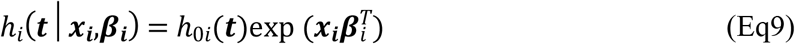

where: *h*_0*i*_(*t*) is the baseline hazard for class *i*; and exp (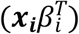) is the relative risk associated with a vector of the ***x_i_*** covariates acting as candidate predictors. We can then derive a survival function from equation Eq9 as follows:

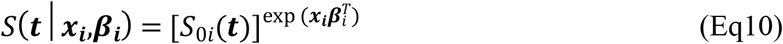

where:

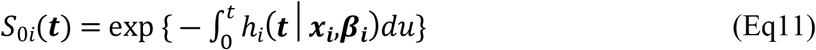

is the baseline survival for individuals at times ***t***, given a vector of candidate predictors ***x_i_*** for class *i*.

The baseline hazard *h*_0*i*_(***t***) in Eq9 is assumed to be an unknown, arbitrary and non-negative function of time and the only parametric part of the model in Eq10 is exp 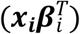 [25]. The maximum likelihood procedure fails to estimate parameters for the likelihood function of Eq9 accurately because the baseline hazard function is not assumed to take any particular form. Instead, these parameters can be estimated using the partial-likelihood approach [26]. This is derived by taking the product of the conditional risk at time *t*_*i*_ given the set of individuals not yet dead, lost to follow-up or censored by that time.

### Part 2. Model specification, selection, validation and software used in Procedures 1, 2a/b, 3a/b and 4

#### Model specification, selection and validation

All subsets regression was deployed [27], along with *k*-fold cross-validation as recommended by Grimm et al. [28], to find the best-fitting model for Procedures 1-4, with four covariates considered for both Cox proportional hazards models *and* (where applicable) the latent class models. The area under the receiver operating characteristic (ROC) curve (AUC) was used to evaluate all models generated – an approach that has been widely used in medical research to assess the diagnostic acuity of biomarkers to discriminate between diseased and healthy subjects [29-31]. In this way the AUC was used in the present study to quantify the extent to which each modelling Procedure was able to discriminate between individuals/participants and classes at risk of mortality. AUC values range from 0.5 to 1, where 0.5 indicates that the discrimination achieved is equivalent to (and no better than that that could be achieved) by chance; a value of 1 indicates perfect discrimination; and a value >0.8 is interpreted as evidence of good discrimination. *k*-fold cross-validation involved randomly dividing the dataset into *k* partitions of approximately equal size, where *k* – 1 partitions were used as a training set and the model was evaluated and validated using the remaining *k*^th^ partition, repeated *k* times. The value *k* = 10 was chosen based on established (and evaluated) best practice [32], with *k* = 10 favoured for less biased model parameters, according to experimentation [33]. The AUC was calculated for each of the 10 test samples, with subsequent confirmation of the results obtained from 10 iterations assessed using a bootstrap re-sampling procedure 1000 times (creating datasets from the original data without making further assumptions) to provide empirical 95% confidence intervals [34].

Covariate selection was guided by the desire to achieve parsimonious models according to the Bayesian Information Criterion (BIC) – the statistic preferred as the most parsimonious penalised likelihood statistic to minimise the risk of overfitting [35]. In choosing the optimum number of latent classes for the latent variable models (i.e. LCA and LCR), BIC was again the preferred statistic as simulations have demonstrated it outperforms other model fit statistics [36]. Strategies for determining the optimal number of classes may also be influence by interpretability (such as clinical salience and/or utility [33,37]), which is often reflected in ‘entropy’ [38] – a measure of the consistency between the modal (highest probability) and probabilistic (exact probabilities) assignment of individuals to latent classes. A high entropy indicates that individuals are more aligned to a single class (large modal probability), which leads to clearer interpretation of each latent class. A low entropy does not preclude latent classes having utility and substantive meaning, but individuals may not be as clearly aligned to just one class, making modal assignment a poor representation of the latent class structure.

#### Software

An important challenge with latent class modelling is its sensitivity to starting values, because these are used to maximize the likelihood function when estimating model parameters. Where the starting values are far from the optimum solution, the likelihood function takes longer to converge and may even fail to do so. Occasionally, up to 50% of the random starts chosen will generate meaningful solutions when the likelihood function is maximized. For a solution to be meaningful, the highest likelihood value is expected to be replicated many times. When this does not occur, it signifies that either: no solution has been achieved and the number of random starts needs to be increased to converge on a global optimum solution; or the specified model structure is unsuitable for the given dataset. While this can add to the time required to explore optimum solutions, once the target values are estimated they can be used as initial values for the final models derived, thereby reducing the duration of the final search process [23].

